# A bench-top dark-root device built with LEGO® bricks enables a non-invasive plant root development analysis in soil conditions mirroring nature

**DOI:** 10.1101/2023.02.12.528178

**Authors:** Georgi Dermendjiev, Madeleine Schnurer, Ethan Stewart, Thomas Nägele, Giada Marino, Dario Leister, Alexandra Thür, Stefan Plott, Jakub Jeż, Verena Ibl

## Abstract

Roots are the hidden parts of plants, anchoring their above ground counterparts in the soil. They are responsible for water and nutrient uptake, as well as for interacting with biotic and abiotic factors in the soil. The root system architecture (RSA) and its plasticity are crucial for resource acquisition and consequently correlate with plant performance, while being highly dependent on the surrounding environment, such as soil properties and therefore environmental conditions.

Thus, especially for crop plants and regarding agricultural challenges, it is essential to perform molecular and phenotypic analyses of the root system under conditions as near as possible to nature (#asnearaspossibletonature). To prevent root illumination during experimental procedures, which would heavily affect root development, dark-root (D-Root) devices (DRDs) have been developed. In this article, we describe the construction and different applications of a sustainable, affordable, flexible, and easy to assemble open-hardware bench-top LEGO® DRD, the DRD-BIBLOX (Brick Black Box).

The DRD-BIBLOX consists of one or more 3D-printed rhizoboxes which can be filled with soil, while still providing root visibility. The rhizoboxes sit in a scaffold of secondhand LEGO® bricks, which allows root development in the dark as well as non-invasive root-tracking with an infrared (IR) camera and an IR light emitting diode (LED) cluster.

Proteomic analyses confirmed significant effects of root illumination on barley root and shoot proteome. Additionally, we confirmed the significant effect of root illumination on barley root and shoot phenotypes. Our data therefore reinforces the importance of the application of field conditions in the lab and the value of our novel device, the DRD-BIBLOX.

We further provide a DRD-BIBLOX application spectrum, spanning from investigating a variety of plant species and soil conditions as well as simulating different environmental conditions and stresses, to proteomic and phenotypic analyses, including early root tracking in the dark.

## 1 Introduction

Plants are sessile organisms, their roots provide anchorage and support for the shoot, and they are key factors regarding the uptake and translocation of water, nutrients as well as the interaction with microbiota (Hodge, 2010; de la Fuente Cantó et al., 2020). Therefore, roots are indispensable when it comes to plant productivity (Lynch, 1995), as they are important for gravitropic response (Žádníková et al., 2015), serve as storage organs and interact with the rhizosphere (Zhu et al., 2011). The root system architecture (RSA) describes the spatial configuration of plant roots in the soil (Smith and De Smet, 2012), which is known to correlate with general crop performance (Zhu et al., 2011). The bio-physicochemical properties of the soil dynamically affect the response of the RSA which is depending on the plant genotype and soil conditions.

With regard to crops, the influence of the RSA on resource acquisition efficiency, plant adaptation to environmental changes and soil-root interactions have been widely studied, since RSA affects crop productivity (de Dorlodot et al., 2007). The transition from germination to subsequent seedling development is initiated by protrusion of the radicle through the coleorhiza, forming the primary root (Weitbrecht et al., 2011). Concomitant with the formation of the coleoptile, seminal roots and crown roots are formed, constituting the majority of the monocot root system. Seminal roots emerge from the primordia in the embryo of the seed, whereas crown roots are post-embryonically formed and emerge from below-ground surface stem nodes (Smith and De Smet, 2012). Interestingly, the angle of growth and the angle between the first appearing seminal roots at the seedling stage are prototypical of the mature RSA in wheat (Oyanagi et al., 1993; Manschadi et al., 2008), and are subsequently considered as representative trait for mature RSA (Richard et al., 2015).

The observation of the root development, or of mature roots, by root phenotyping is especially important for the identification of root traits for crops and finally for crop yield improvement (reviewed in Tracy et al., 2020). Additionally, huge effort is put into studying RSA of the conventional dicot model plant *Arabidopsis thaliana*, where *in vitro* studies on agar plates (Xiao and Zhang, 2020) as well as *in situ* studies in rhizotrons have been performed (Rellán-Álvarez et al., 2015; Ogura et al., 2019; LaRue et al., 2022). Because of image acquisition setups via cameras or scanners, roots are often exposed to light, which heavily affects the root development (Cabrera et al., 2022). Recent studies in *Arabidopsis thaliana* emphasize even the negative effect of light on root development, and scientists consequently shift to implementing DRDs in their approaches (Silva-Navas et al., 2015; Silva-Navas et al., 2016; Garcia-Gonzalez et al., 2021). Thus, to get meaningful results that are applicable to the field, it is indispensable to analyze the root architecture in the dark in soil conditions mirroring field conditions, especially for crop root phenotyping.

Rhizoboxes have been used for two-dimensional (2D) root visualization since the 1980s (Marschner and Römheld, 1983; Youssef and chino, 1987; Fitter et al., 1988). In 2D approaches, compared to a possible three-dimensional (3D) root development in pots, roots are forced to grow in 2D along a (glass) slide (Nagel et al., 2015; Bodner et al., 2017) due to angled rhizoboxes and gravitropism. Additionally, since phenotypes in shoots and roots are expressed differently depending on the soil conditions, including soil water content and temperature, whole-plant phenotyping is emphasized, where roots and shoots are measured simultaneously (reviewed in Tracy et al., 2020). Thus, rhizoboxes are an optimal way to analyze root growth development in parallel to shoot development without any effect of the root:shoot ratio (Mašková and Klimeš, 2020).

Currently, the setup of lab experiments is challenging. The COVID-19 pandemic showed us the dependency on an efficient supply pipeline since the scientific output was impacted due to lack of lab supplies (Heo et al., 2022). Additionally, a great effort is made by scientists to reduce the anthropogenic climate change, by including more sustainable research.

Inspired by these different challenges, we established a non-invasive, sustainable bench-top DRD that enables whole-plant molecular analysis and phenotyping in conditions as near as possible to nature (#asnearaspossibletonature). We used predominantly secondhand materials (LEGO® bricks), materials produced in our lab, already available resources, or we bought locally to reduce the CO2 footprint. We chose LEGO® bricks to build a dark housing for the rhizoboxes that is flexible in size, resistant to environmental parameters, and easily transportable. Originally used as toy, LEGO® bricks have already inspired a variety of teachers and scientists to translate knowledge and to use these bricks for scientific applications (Lin et al., 2018; Mäntylä and Ihalainen, 2021; Montes et al., 2021). In plant science, LEGO® bricks have been recently used for building small-scale engineered environments for plant roots (Lind et al., 2014).

The LEGO® brick DRD-BIBLOX, short BIBLOX, can house between one and fourteen in-house made rhizoboxes in a small setup. We show, that the BIBLOX can be used for a wide application range, including whole-plant proteomic analysis and root phenotyping of crops grown in different soil compositions mirroring natural field conditions. We also include stress applications and the analysis of the root growth and morphology over real time. The BIBLOX is especially applicable for the analysis of molecular biology-related investigations (e.g. reverse and forward genetic approaches) and for analyses of the RSA in response to distinct environmental factors including different substrate composition.

## 2 Material and Methods

### Monitoring the environmental parameters enables the translation of natural environmental conditions to controlled lab conditions

Soil temperature, soil water content, light intensity and air humidity were measured in a barley field (9 ha) in Lower Austria in the years 2021 and 2022. For the measurements in the year 2021, a TensioMark® sensor (ecoTech Umwelt-Messsystem GmbH, Bonn, Germany) was used to measure the soil moisture (pF-value) and the soil temperature. Three sensors run by one data logger each were positioned within the 9 ha field, with 60 meters between measuring points. The sensors were mounted at -30 cm soil depth. The data was saved using the Data logger “envilog Maxi” (ecoTech Umwelt-Messsystem GmbH, Bonn, Germany). The environmental parameters measured in the year 2021 are published at our homepage www.celbics.com. In the year 2022, we used three TekBox-TBSST04-3 (TR) temperature measuring sensors (Umweltanalytische Produkte GmbH, Cottbus, Germany) accumulating the data in soil depth of -20, -30, and -50 cm and three PR2/4 SDI-12 Delta-T profile sensors (Umweltanalytische Produkte GmbH, Cottbus, Germany) accumulating the data in -10, -20, - 30, and -40 cm of soil depth, respectively (**Figure S1A, Supplemental Video 1**). The data was saved by one solar powered Datalogger (yDoc ML-317, Firmware version 4.3 build 8) (Umweltanalytische Produkte GmbH, Cottbus, Germany). Three Apogee quantum sensors Model SQ-421 were used to measure the photosynthetically active radiation (PAR) in the field. Measurements were performed every 15 minutes and saved to an SD-Card. After the end of the field trial, the data was exported into .csv format via the Software “ydocTerminal” version 3.13. The air temperature was measured with three Lascar EasyLog data loggers (Lascar electronics, Wiltshire, United Kingdom), each positioned in a weather house mounted on a wood pole in 80 cm hight (**Figure S1A**).The data was imported into “RStudio” version 2022.12.0+353 (RStudio Team, 2020) with “R core” version 4.2.2 (R Core Team, 2022). Rows with missing values were filtered out, as well as soil moisture sensor 3, which delivered only very few datapoints (probably due to voltage drop in the cable) and one of the soil temperature sensors at -20 cm depth, which got damaged during the setup and sent incorrect data. Air temperature and humidity data was imported from the three sensors and merged. Data was subset for the first 16 days, the wanted sensor (and depth for soil moisture- and temperature) and transformed into long format using the “melt” function from the R-package “reshape2” (Wickham, 2007). Plots were created using the R-package “ggplot2” (Wickham, 2016) with the color palette “Set2” from the R package “RColorBrewer” version 1.3-3 (Neuwirth, 2022). PAR and soil temperature datapoints aligned almost perfectly for the replicates and were therefore only drawn using the “geom_line” function. Datapoints of the soil moisture and air temperature showed greater variability, therefore smoothened means of the datapoints were plotted using the function “geom_smooth” with the parameters: method = “loess” and a span of 0.1 for the soil moisture and 0.01 for air temperature **(Supplemental Data1)**.

### Construction of the BIBLOX

#### Material for DRD-BIBLOX

.) 3D-CAD software Fusion 360 (Autodesk Inc, San Rafael, California, USA)
.) 3D printer Ultimaker S5
.) Polylactic acid (PLA) filament for 3D printing
.) Rhizoboxes, 200 mm x 150 mm x 30 mm
.) LEGO® DRD-BIBLOX (composed of around 800 black LEGO® bricks and 80 plates)
.) Infra-red (IR) LED Cluster_880 nm 5 mm T-1 3/4 (Kingbright, BL0106-15-29)
.) Glass (2 mm)
.) Plant growth chamber (Conviron)
.) Raspberry Pi3 B+ single board computer
.) Pi3 Camera (Electreeks® Raspberry Pi camera module with an automatic infrared cut filter - full HD
.) 75.5° standard tripod
.) Foam rubber, 1.7 mm thickness
.) ImageJ Software (https://imagej.nih.gov)
.) The R Project for Statistical Computing

#### Rhizobox setup and design

Initial rhizoboxes were constructed home-made from 5 mm polyvinyl chloride (PVC) sheet and bonded with silicon-based glue (**Figure 1A**). Additionally, 3D-printed versions were designed using the 3D-CAD software Fusion 360 **(Supplemental Data2)**. Rhizoboxes were printed using an Ultimaker S5 3D with PLA filament (**Figure 1B**). Dimensions of both versions were 200 mm height, 150 mm width, and 30 mm depth.

**Figure 1.**
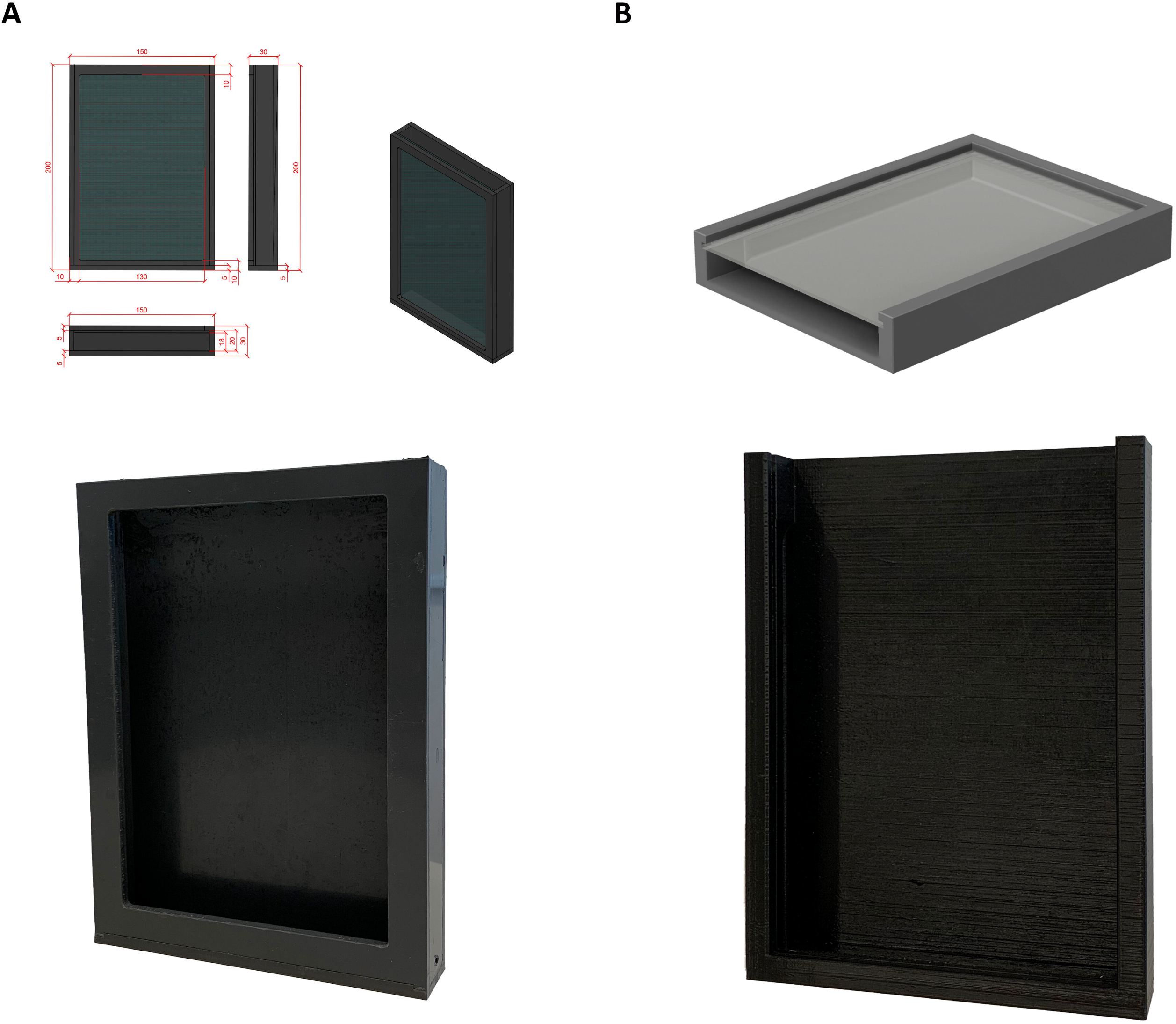
Rhizoboxes constructed from PVC sheet (A) and 3D printed (B).

#### Set up of the BIBLOX

Black secondhand LEGO® bricks were used to construct the base of the BIBLOX. Special LEGO® bricks and plates, which were not available pre-owned were bought locally. Only a few rare bricks were bought online from LEGO® A/S (https://www.lego.com/en-gb). The rhizobox was positioned at an angle of 60 degrees within the BIBLOX by blades attached to the inner brick walls.

#### Image acquisition

The construction of the BIBLOX took around 60 minutes (**Supplemental Video 2**). Once the BIBLOX was constructed, we installed the Pi3 camera and the IR LED cluster within the box. The Pi3 camera was mounted on a 75.5° standard tripod and connected to the Raspberry Pi3 B^+^ computer placed outside the box. The computer was connected to LAN network (Wi-Fi would also be possible). The camera was set to capture an image every four minutes. The IR LED Cluster source was controlled by a relay (Raspberry Pi Power Relay Board Expansion Board Module Three Channel (3-ch)) installed on the Raspberry Pi3 computer, which switched the light source on for only one second during image acquisition. In this way, we limited root light exposure to a minimum.

The BIBLOX inclusive the set-up for non-invasive root tracking costs around 520€.

### Plant materials and growth condition

The spring barley (*Hordeum vulgare L*.) wild-type variety Golden Promise (GP) and the facultative variety BCC93 (kindly provided by Kerstin Neumann, IPK Gaterseleben) were grown in the plant growth chamber (PSI) at 14 °C/12 °C, maize and wheat (Bobwhite) at 16 °C/14 °C, 12 h day/12 h night cycle with light intensity of 130 - 220 µmol m^-2^ s^-1^ and 70% humidity. To track early root growth, GP and BCC93 were grown #asnearaspossibletonature, according to (Dermendjiev et al., 2021) and the measured data of the field experiments, at 14 °C/12 °C 12 h day/12 h night cycle with light intensity of 130 - 220 µmol m^-2^ s^-1^ and 70% humidity, in a plant growth chamber (Conviron Adaptis A1000). Tomato was grown in the glass house at 26 °C/19 °C, 12 h day/12 h night cycle with light intensity between 650 and 4 000 Lux (lx) and between 50 – 60% humidity.

### Soil compositions

Within this project, we used five different soils: (I) For the growth of tomato, maize and wheat, we used sieved (3 mm mesh) Cocopeat (CP) supplied with H2O. (II) To enable a high root highlighting for the imaging, we used sieved (3 mm mesh) CP that was mixed with activated carbon 2:1, supplied with H2O (CP*). Peat substrate (Gramoflor) was supplied with H_2_O. We used naturally grown bio-organic field soil (BFS) (III) and naturally grown conventional field soil (CFS) (IV) that were obtained from a field in Lower Austria. BFS and CFS were analyzed by the Austrian Agency for Health and Food Safety GmbH, in short AGES (Vienna, Austria). BFS shows a higher pH-value (pH = 6.3) and less mineral nitrogen (0.3 mg/100 g) compared to CFS (pH = 5.5; mineral nitrogen: 0.5 mg/100 g) (**Supplemental Data 3, 4**). For salt stress analysis, we mixed CP* with water containing 20 g NaCl/l H_2_O (electric conductivity (EC) = 30 EC), CP*_30EC (V). Soils were adjusted to a pF value 2-3 to enable a water moisture content mirroring soil environmental parameters #asnearaspossibletonature and were added to the rhizoboxes, respectively.

### Sampling for proteomic analyses

For proteomic analyses root and shoot material of GP was used. Barley grains were germinated in rhizoboxes filled with soil (CP*). Seven grains were sowed per rhizobox. For control conditions rhizoboxes were inserted into a BIBLOX setup in a climate chamber, therefore roots would develop in dark. For root illumination conditions stand-alone rhizoboxes were put directly in a climate chamber without covering the roots in the rhizobox. All rhizoboxes were installed at an angle of 60 degrees allowing root growth along the glass front of the boxes. Conditions in climate chambers (Conviron Adapsis CMP 6010, Controlled Environment limited) were set to 12-hour day/night cycles of 14/12 °C, with a light intensity of 130-220 µmolm^-2^s^-1^ and 70% humidity. No difference in the soil temperature was measured between the stand-alone rhizoboxes and the rhizoboxes covered by the BIBLOX. Seedlings were harvested 8 and 16 days after sowing (DAS) respectively. Root (R) and shoot (S) material was separately harvested, cleaned from soil, and immediately frozen in liquid nitrogen (LN_2_). Shoots and roots of plants with light grown roots (LGR) were harvested under light conditions (S_LGR + LGR). And plants of dark grown roots (DGR), the shoot was harvested in light and the root was harvested in a completely dark room with dimmed red light to prevent root illumination (S_DGR + DGR). Combined root or shoot material of 14 plants for 8 DAS and seven plants for 16 DAS would count as one biological replicate respectively. Three to four biological replicates were taken per category (8 DAS_S_DGR, 8 DAS_S_LGR, 8 DAS_DGR, 8 DAS_LGR, 16 DAS_S_DGR, 16 DAS_S_LGR, 16 DAS_DGR, 16 DAS_LGR).

### Protein extraction and digestion

Material was homogenized to powder, using LN2, mortar and pestle. Proteins were extracted using a Sucrose SDS-buffer [100 mM Tris-HCl pH 8.0, 30% (w/v) Sucrose, 0.5% (v/v) 2-Mercaptoethanol, 10 mM EDTA, 2% (w/v) SDS, Protease Inhibitor (Roche, Cat. No. 05 892 791 001)] by adding 1 ml of buffer to 350 mg of sample. Samples were resuspended completely. 750 µl ROTI®Phenol [Roth, Cat. No. 0038.3] were added to the samples for protein extraction. Samples were vortexed for 1 min and incubated for 5 min followed by centrifugation at 20,000×g for 5 min at room temperature (RT). After phase separation, the phenol phase was carefully transferred to a new tube. The phenol fractions were counter-extracted with 750 µl of Sucrose SDS-buffer, vortexed for 1 minute and incubated for 5 min and then centrifuged at 20,000×g for 5 min at RT. The phenol phase was carefully transferred to a new reaction tube. Proteins were precipitated by adding 2.5 volumes of ammonium acetate in methanol [0.1 M]. After 16 hours incubation at -20°C, proteins were pelleted by centrifugation at 4°C for 5 min at 5,000×g. Supernatants were discarded and the protein pellets were washed with ice cold ammonium acetate in methanol [0.1 M] and 70% methanol respectively, followed each by centrifugation at 4°C for 2 min at 18,000xg. The supernatant was removed, and protein pellets were air dried for 60 min and subsequently resuspended in 50µl urea buffer [8 M urea, 100 mM ammonium bicarbonate, 5 mM DTT, Protease Inhibitor] while incubated at 37°C for 20 min for better solubility. Next samples were centrifuged at RT at 20,000xg for 10 min.

Protein concentration was measured via Bradford assay using a Quick Start™ Bradford 1x Dye Reagent (Biorad, Cat. No. 5000205) prior to protein content normalization. BSA (Bovine Serum Albumin) dilution series (0 – 10 mg/ml) in the according buffer were used as standard to calculate sample protein concentration. 2 µl of sample or standard were pipetted into 1.5 ml tubes (in triplicates). 1 ml of Bio-Rad Quick Start™ Bradford 1x Dye Reagent was added. Tubes were vortexed and incubated in dark for 10 min. 200 µl of the solution were transferred into a 96-well plate. The absorbance of standards and samples was measured at 595 nm wavelength using a Thermo Scientific Multiskan Spectrum. BSA standard curve and calculation of protein concentration were done using Microsoft Excel. Cystein residues were reduced by incubating 200 μg protein per sample for 45 min at 30°C while shaking at 700 rpm. Cysteine residues were alkylated with 55 mM Iodoacetamide (IAA) while shaking with 700 rpm, in the dark, at RT, for 60 min. Increased DTT [10 mM] concentration and sample incubation at RT, shaking at 700 rpm for 15 min stopped alkylation process.

Further the urea concentration was diluted to 2 M with 100 mM ammonium bicarbonate/10% Acetonitrile (ACN). CaCl2 was added to a final concentration of 2 mM. Trypsin digestion was performed at 37 °C rotating for 14-16 hours using Poroszyme™ Immobilized Trypsin Cartridge (ThermoScientific Cat. No. 8-0087-40-0994) at a ratio of 5:100 v:w. Peptides were desalted using C18 solid phase extraction columns (Bond Elut SPEC C18, 96 round-well plate, 15 mg, 1 mL, Agilent Technologies, Santa Clara, USA) and a water-jet (vacuum) pump. Plates were activated with 2×400 µl methanol passing the columns by gravity for 2 min and then aspirated via the pump. Columns were equilibrated with 4 × 400 µl of ultrapure H2O, passing the column by gravity for 2 min and then aspirated via the pump. Subsequently samples were pipetted into column and peptides and salt would bind to it while gravity flow for 5 min, followed by aspiration via pump. Subsequently samples were desalted with 5 × 400 µl of ultrapure H2O passing the column by gravity for 2 min and then aspirated via the pump, last aspiration to total dryness. Purified peptides were recovered with 2×200 µl of Methanol, passing the column by gravity for 5 min and then total aspirated via the pump. Peptides were transferred into new tubes and dried in a SCANVAC CoolSafe Vacuum Concentrator for 5 hours at RT. The peptides were resuspended in 0.1% Formic Acid (FA) in acetonitrile. The final peptide concentration was measured spectrophotometrically via a NanoDrop device (Thermo Scientific).

### LC-MS/MS analysis

Liquid Chromatography and Tandem Mass Spectrometry analysis was performed on a nano-LC-system (Ultimate 3000 RSLC; Thermo Fisher Scientific) coupled to an Impact II high resolution quadrupole time-of-flight (Bruker) using a Captive Spray nano electrospray ionization source (Bruker Daltonics). The nano-LC system was equipped with an Acclaim Pepmap nanotrap column (C18, 100 Å, 100 µm 2 cm; Thermo Fisher Scientific) and an Acclaim Pepmap RSLC analytical column (C18, 100 Å, 75 µm × 50 cm; Thermo Fisher Scientific). The peptide mixture was fractionated by applying a linear gradient of 5% to 30% solvent B [0.1% FA in acetonitrile] at a flow rate of 250 nl min^−1^ over a period of 60 min, followed by a linear increase 30-45 % solvent B within 15 min. The column temperature was set to 50°C. MS1 spectra were acquired at 3 Hz with a mass range from *m*/*z* 200–2000, with the Top-18 most intense peaks selected for MS/MS analysis using an intensity-dependent spectra acquisition time between 4 and 16 Hz. Dynamic exclusion duration was 0.5 min.

### Data analysis and visualization

Proteomics MS raw files were processed using the MaxQuant software (version 2.0.30; Cox and Mann, 2008). Peak lists were compared against the barley reference proteome (*Hordeum vulgare subsp. vulgare* (Domesticated barley), cv. Morex, Uniprot, UP000011116, version March 2022) using the built-in Andromeda search engine (Cox et al., 2011). Enzyme specificity was set to trypsin, allowing up to two missed cleavages. Cysteine carbamidomethylation was set as static modification, and N-terminal acetylation and methionine oxidation as variable modifications. During the search, sequences of 248 common contaminant proteins and decoy sequences were automatically added. A false discovery rate (FDR) of 1% was set at peptide and protein level. Proteins were quantified across samples using the label-free quantification (LFQ) algorithm (Cox et al., 2014) and the match-between-runs option was enabled.

Uncharacterized proteins were manually identified by using the Uniprot BLAST application. Data was analyzed and visualized using Microsoft Excel (version 2211 Build 16.0.15831.20098 for Microsoft 365 MSO), and RStudio (version 2022.02.2 for Windows). Proteins of which LFQ values were not detected for any of the measured sample groups as well as proteins where three out of four or two out of three values of biological replicates of one sample group were missing were dismissed. For missing third (for three replicates) or forth (for four replicates) values and average value was calculated from the other two of three values of the biological replicates (**Supplemental Data 5**).

The T.TEST function (heteroscedastic, with two tailed distribution; Microsoft Excel) was used to find significant differences regarding LFQ values between mean values of different sample groups (**Supplemental Data 5**). Prior to PCA (principal component analysis) data was logarithmically normalized using log10(x+1). PCAs, Loading plots and Contribution plots of all data and subgroups were calculated and visualized using RStudio (**Supplemental Data 6, 7)**. Proteins that significantly differed in their abundance when comparing plants (shoots and roots) of light growing roots (LGR) and dark growing roots (DGR) were classified regarding subcellular localization and molecular function using Microsoft Excel. Subcellular localization categories were “Cytoplasm”, “Cytosol”, “Nucleus”, “Mitochondria”, “integral component of membrane”, “Ribosome”, “Chloroplast”, “Extracellular”, “Plasma membrane”, “Cytoskeleton”, “Endoplasmic reticulum”, “Golgi apparatus”, “Peroxisome”, “Vacuole”, “Cell wall”, “Apoplast” and “Plasmodesmata”. And molecular function categories included among others “RNA binding”, “ATP binding”, “metal ion binding”, “Oxidoreductase activity”, “defense response activity”, “Cytoskeleton” and “Actin filament binding” (**Supplemental Data 5**).

### Image analysis

To enhance the contrast between the roots and substrate for further semi-automatic image analyses, we manually traced the roots in every 10^th^ image from each experiment (derived from between three and five biological replicates) with an Apple Pencil on an iPad (**Figure S2A, B**), since those tools have already been available in our lab as we use digital lab notebooks to work paperless in our group. From the traced images, a binary image of the root system was made using colour thresholding. Binary images were skeletonized and a network graph was constructed using the sknw package (Xiaolong, 2020). From the network graph, the longest root and total root system length was calculated using the network-x package (Hagberg et al., 2008). Primary root angle was calculated by fitting a line through the x and y coordinates of the primary root skeleton pixels. The convex hull area and bounding box width of the root system was calculated from the binary images using OpenCV (Bradski, 2000) (**Figure S2 C**).

The root growth angle (RGA) and the seminal root growth angle (SRGA) was measured with the angle tool provided in ImageJ (Schneider et al., 2012) by drawing lines from the grain to the maximum distance of the seminal root to the horizontal level of the grain and between the first two seminal roots (**Figure S2 D**). Data was analyzed and visualized using GraphPad Prism (version 9.0 for Mac, GraphPad Software, San Diego, California USA, http://www.graphpad.com/).

## 3 Results

### 3.1 Monitoring the environmental parameters enables lab experiments as near as possible to nature (#asnearaspossibletonature)

Recently, we have successfully set up conditions to follow the germination in the lab at parameters #asnearaspossibletonature (Dermendjiev et al., 2021). We monitored the soil temperature and moisture, air temperature and PAR in a field of an organic spring barley farmer in lower Austria between the period of sowing and harvesting barley within 2021 and 2022, respectively. The measured data of 2021, which are publicly available at our group homepage (www.celbics.com) show a pF-value in -30 cm depth for the first 16 days between 2 and 2.5. Additionally, the soil temperature was between 6 – and 13 °C in -30 cm depth. In 2022, the soil temperature was between 7.3 and 18.2 °C in -20 cm soil depth resulting in a mean temperature of 12 °C (**Figure S1 A, B**). The soil water content was between 14 and 20 % (v/v). Thus, the parameters measured in year 2022 were consistent with the environmental parameters measure in the year 2021. Subsequently, the temperature for barley germination in the lab condition was set to the temperature 14 °C/12 °C, considering that barley grains are sown at -3 cm below the soil surface and that air temperature was measured between 0 °C and 25 °C within these first 16 days. The soil moisture was set for the germination to a pF-value between 2-3 during the first 16 days for germination.

### 3.2 The construction of rhizoboxes for non-invasive *in situ* early root tracking: from home-made PVC constructed rhizoboxes to 3D printed version

Rhizoboxes were constructed from PVC for the purpose of using them as DRD but also as stand-alone devices. To reduce the CO2 footprint and increase flexibility, subsequently rhizoboxes were 3D printed using polylactic acid (PLA), a thermoplastic polymer which is manufactured from renewable and biodegradable plant-based materials (Henton et al., 2005; Bhatia et al., 2007) (**Figure 1**). Additionally, PLA is able to withstand plant growth conditions. This 3D printed version enables high flexibility in terms of construction size and timepoint. Additionally, using 3D printers is a first small step for more sustainable research in the lab.

### 3.3 The diverse applications of the BIBLOX for non-invasive *in situ* early root development analysis

Applying the measured natural environment parameters enables us to perform experiments in controlled lab conditions with settings #asnearaspossibletonature.

#### 3.3.1 BIBLOX allows proteomic analysis of roots and shoots of plants grown #asnearaspossibletonature

As root illumination heavily affects root development, we wanted to provide a device that allows phenotypical and molecular analyses of root and shoot material grown under parameters close to field conditions.

The BIBLOX provides a dark housing for the rhizoboxes that is flexible in size, resistant to environmental parameters, and easily transportable (**Figure 2A, B**). We built the BIBLOX which in our setup covers twelve rhizoboxes for proteomic analysis of shoot and root (**Figure 2B**). To prove the applicability of our device and to reinforce the importance of working as near as possible to nature we performed phenotypic and proteomic analysis of root and shoot material of plants of LGR compared to plants of DGR grown in our BIBLOX which allows root development in darkness. For this approach rhizoboxes were filled with soil and 7 barley grains per rhizobox were put for germination. The BIBLOX with rhizoboxes was put into the growth chamber for up to 16 days. Assessment of the incoming light to the BIBLOX showed a 92% - 95% reduction in light intensity measured at root level (< 10 µmol m ^2^ s^-1^ in the BIBLOX compared to 130 - 220 µmol m^-2^ s^-1^ in the Conviron). Additionally, stand-alone rhizoboxes, positioned at an angle of 60 degrees and without coverage were installed in the climate chamber too, allowing root illumination of developing plants. Pictures were taken at 4, 6, 8 and 16 days after sowing (DAS). A delay in barley root and shoot development in plants of LGR compared to plants of DGR roots could be observed (**Figure 3A**). After 16 days plants were removed from the rhizoboxes, the roots were washed, and images were taken to assess the final root and shoot growth size (**Figure 3B**). DGR were slightly smaller compared to LGR (**Figure 3A, B**). Interestingly, regarding phenotypes, the shoot was much more affected since the shoot length of the plants of DRG were significantly reduced compared to the plants of LGR (**Figure 3B**).

**Figure 2.**
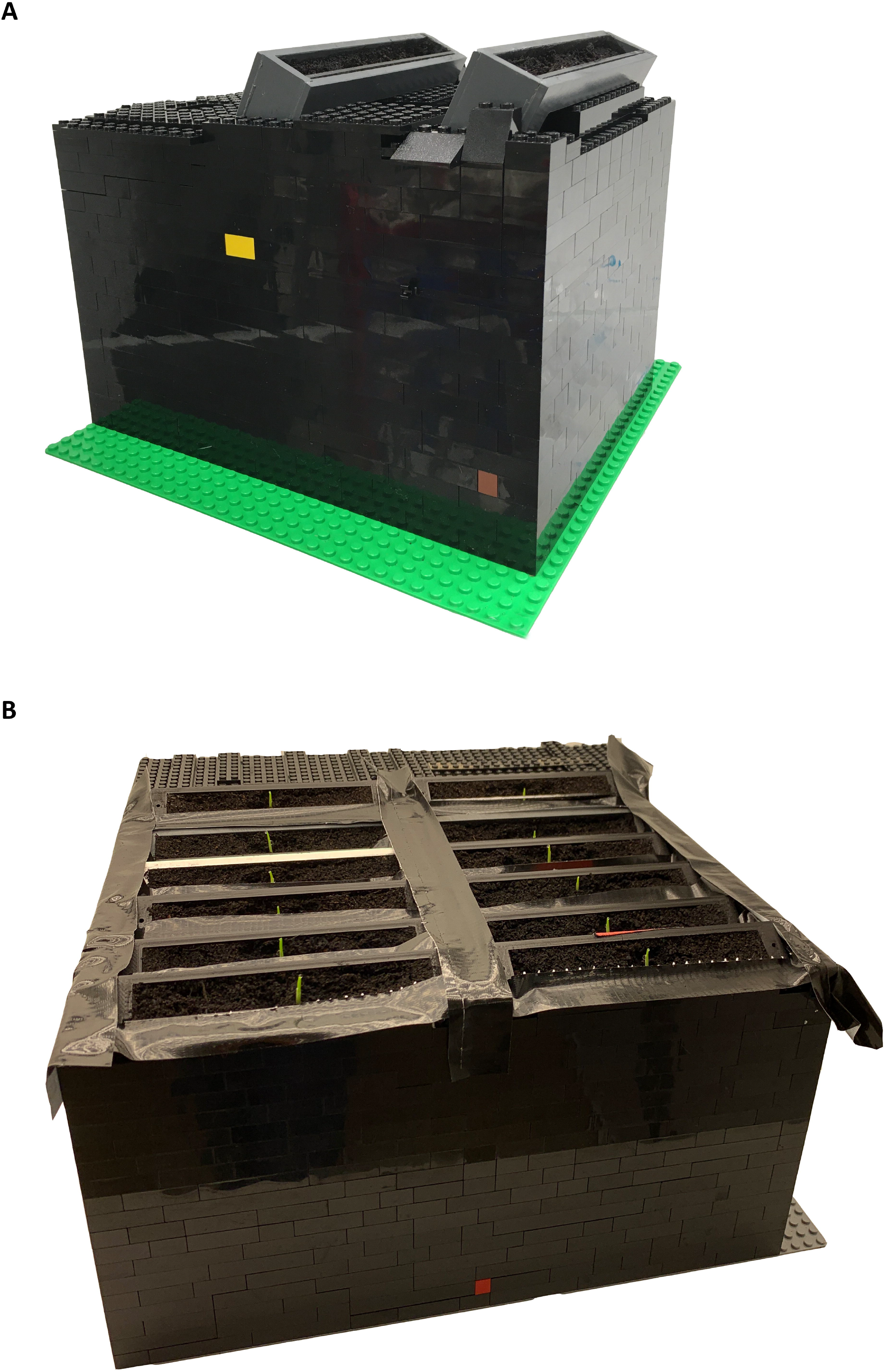
The BIBLOX as housing for (A) two and for (B) 12 rhizoboxes.

**Figure 3.**
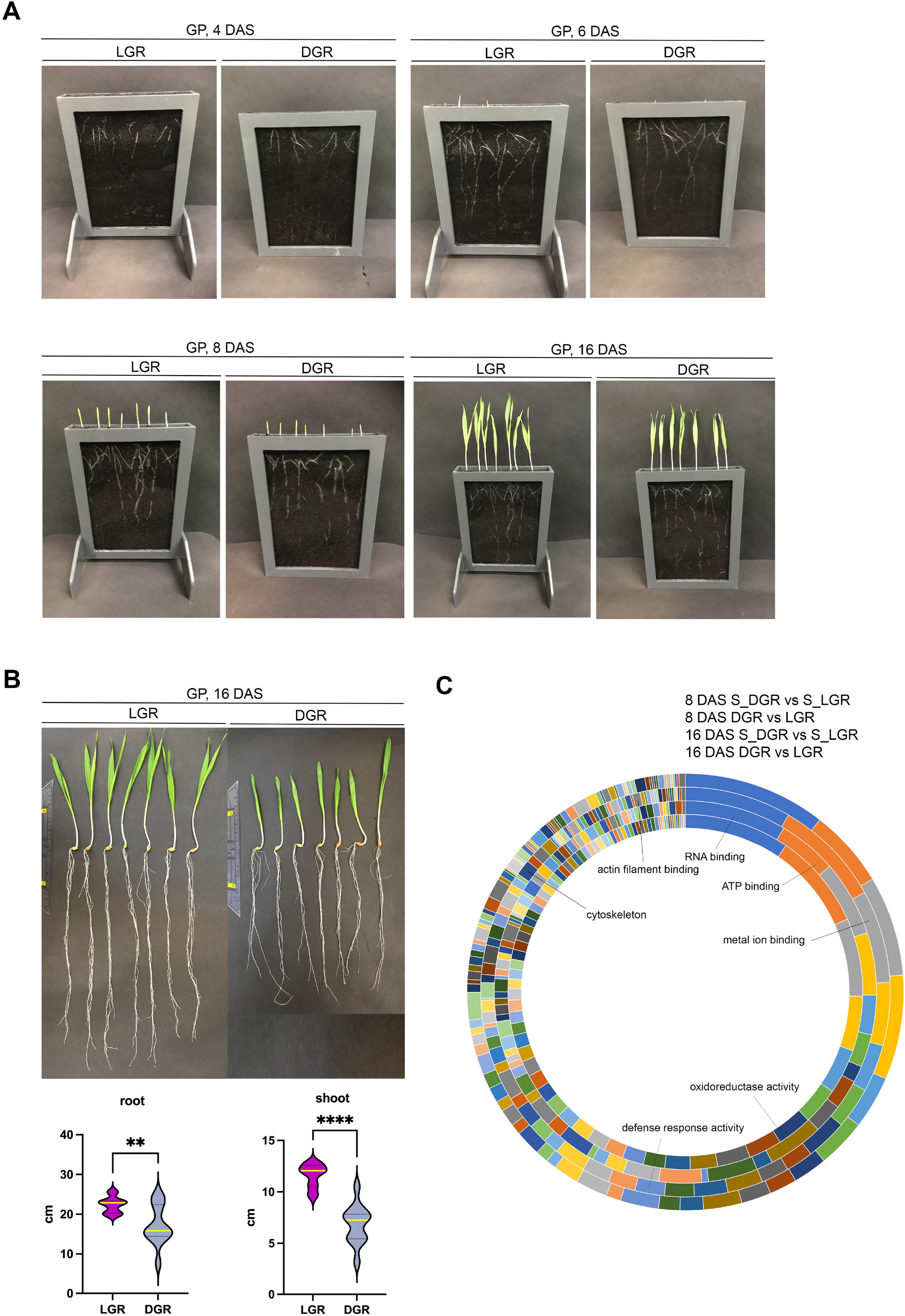
GP growth and proteomic analysis of roots and shoots. (**A**) Pictures of GP seedlings in rhizoboxes grown in CP* were taken at 4, 6, 8 and 16 DAS, respectively. 7 grains were sown in each rhizobox. (**B**) Root and shoot length of plants grown for 16 DAS. Plants were harvested from the rhizobox and washed. Light grown roots (LGR), dark grown roots (DGR). All shoots were in light. Violin plots show the root and shoot size of roots grown in the light (n = 12) and dark (n = 12). ** represents ≤ 0.005, **** represents ≤0.0001. Yellow line represents the median. (**C**) Functional classification of significantly up-or downregulated proteins in root and shoot upon root illumination. Indicated proteins are involved in RNA-, ATP-, and metal ion binding, in the oxidoreductase and defense activity, and cytoskeleton related proteins.

For proteomic sampling plant roots were kept continuously in dark or light for 8 and 16 days. Finally, 8 and 16 DAS roots and shoots were harvested according to their growth light settings. This was followed by protein extraction and digestion and subsequent proteomic analysis. For the root and shoot material, in total, we identified 2158 proteins, 1236 of them show significant changes in their abundance in root and shoot following root illumination. Out of all significantly different regulated proteins upon root light exposure, in 8 DAS roots, about 50% (68% for 16 DAS) where downregulated and 50% (32% for 16 DAS) upregulated compared to dark conditions. While in 8 DAS shoots of plants of LGR, about 47% (38% for 16 DAS) where downregulated and 53% (62% for 16 DAS) upregulated comparing DGR. Principal component analysis (PCAs) of 8 DAS, 16 DAS and of all data showed a clear separation between sample groups and clustering of biological replicates within sample groups **(Figures S3, 4, 5)**. Principal Component 1 (PC1) separates the proteins regarding root and shoot-specificity. PC2 separates the proteins according to root illumination (**Figures S3, 4)**. The PCA of 8 DAS data shows that proteins of LGR and the corresponding shoots are clearly separated from proteins of DRG. Additionally, at 8 DAS proteins of roots and shoots are distinctively separated, too. At 16 DAS, proteins of roots and shoots are clearly separated as well. However, proteins of LGR show only specific separation in roots, but not in the corresponding shoots (**Figures S3, 4)**. According to the PCA plots, the effect of root illumination on the tissue specific protein abundance at 16 DAS appears stronger in the roots compared to the effect in shoots (**Figures S3, 4, 5)**. The subcellular classification of significantly up- or downregulated proteins in root and shoot upon root illumination showed a broad range of protein localizations (**Supplemental Data 5**). Further a classification of molecular functions of those proteins showed them being highly involved in RNA binding, ATP binding, metal ion binding, as well as in oxidoreductase activities, defense response activities, the cytoskeleton and actin filament binding **(Figure 3C)**. Additionally, we found differently regulated protein levels of for example reactive oxygen species (ROS) associated proteins and auxin pathway associated proteins as well as defense response associated proteins and cytoskeleton related proteins upon root light exposure.

These data show that the BIBLOX can be used as an effective DRD for proteomic analysis, since our proteomic data confirms already published effects of light on roots, e.g. on cytoskeleton proteins (Dyachok et al., 2011; Du et al., 2020; Halat et al., 2020; Cabrera et al., 2022) and on ROS (Yokawa et al., 2011). Additionally, first steps into whole-plant phenotyping show the effect of light root illumination on shoot development. Our data therefore reinforces the importance of the application of field conditions in the lab and the value of our novel device, the BIBLOX.

#### 3.3.2 BIBLOX as system for dark-root growth analysis of several crop plants and different soil conditions

Aiming to establish a DRD with a broad application, we further evaluated the application of BIBLOX for growth analysis for additional crops. Tomato, maize and wheat were grown in CP under appropriate settings. The RSAs of the used crops could be clearly observed at 8 and 16 DAS (**Figures 4A)**

**Figure 4.**
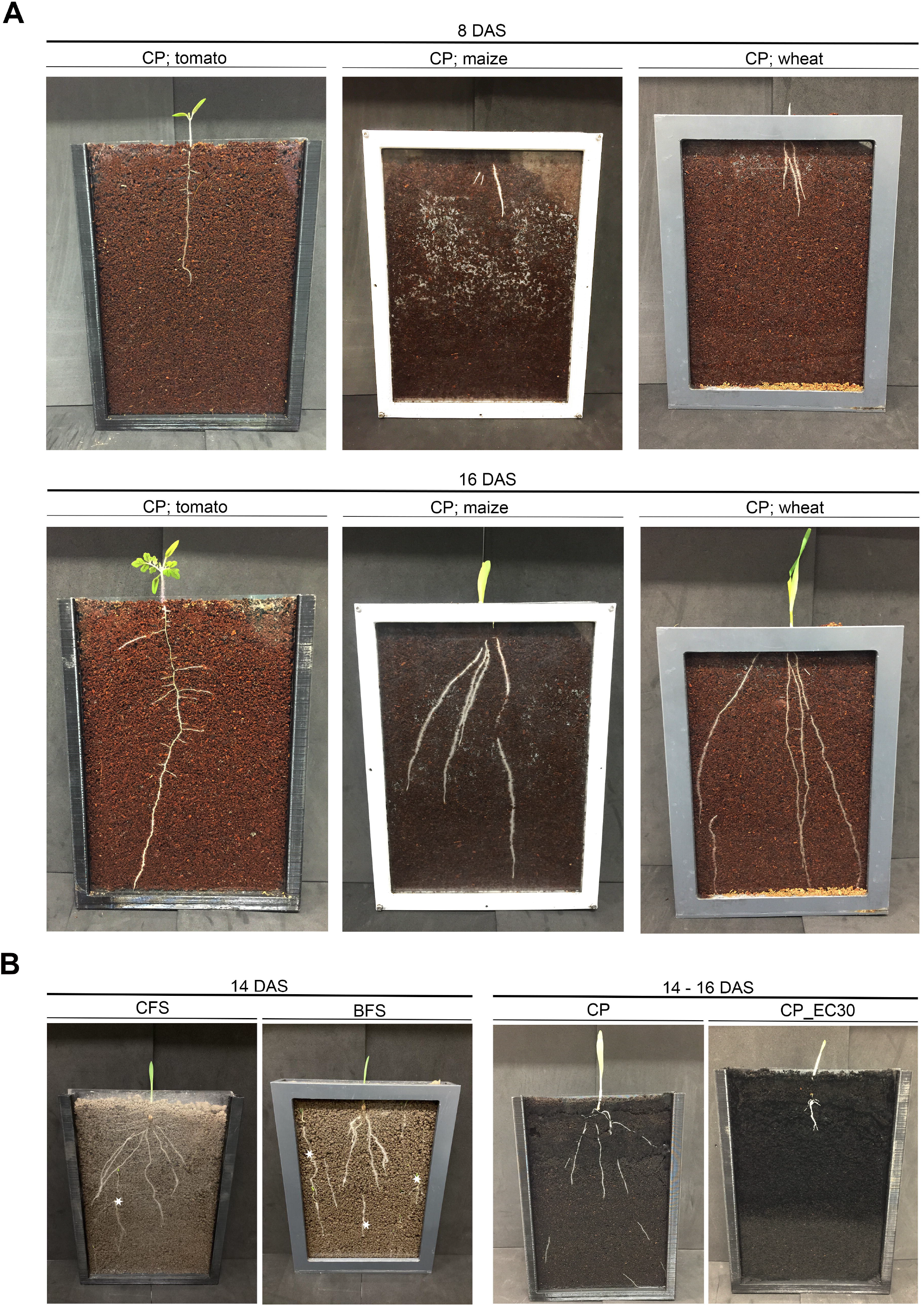
The BIBLOX enables the root growth analysis of distinct crops and the application of nature environmental conditions. (**A**) Plant growth of tomato, maize, and wheat, in CP soil. Pictures were done at 8 and 16 DAS. (**B**) GP was sown in CFS and BFS. A picture was made at 14 DAS. Note the appearance of weeds in the natural field soil, indicated with *. GP was sown in CP*_30 EC. A picture was made at 14 – 16 DAS.

Since non-natural soil conditions alter the root development, the next step with respect to accurate controlled lab experiments is the application of natural soil conditions. Subsequently, we applied naturally grown BFS and CFS and analyzed the root development of GP grown in the BIBLOX. 14 DAS RSA was clearly different from roots grown in BFS compared to CFS (**Figure 4B**). Additionally, GP was exposed to salt stress (30EC) during germination and early root development that corresponds to salt-tolerant conditions that barley, as salt tolerant plant, is able to handle (**Figure 4B**). These data show the diverse application of the BIBLOX to study the RSA of different crops and different soil conditions.

#### 3.3.3 BIBLOX as a system for uninterrupted root growth and morphology analysis over time

To avoid root illumination during root development, we set up a non-invasive root tracking method that enables an uninterrupted root growth. We built the BIBLOX which covers one rhizobox, a light source and the camera for early *in situ* root tracking (**Figure 5**). We used around 800 black LEGO® bricks for the base (**Figure 5A**), around 80 plates (**Figure 5B**), one base plate and eight special pieces for the two holders of the rhizobox (**Figure 5C**). In total, the BIBLOX includes 22 rows of LEGO® bricks (**Figure 5D**), where the holders were placed from the 6^th^ to the 16^th^ row. For the independent biological replicates, we used one BIBLOX for the image analysis. However, upscaling the system is possible with up to three parallel BIBLOXes per shelf within our CONVIRON plant growth chamber (**Figure S6**). Four to seven biological replicates were analyzed.

**Figure 5.**
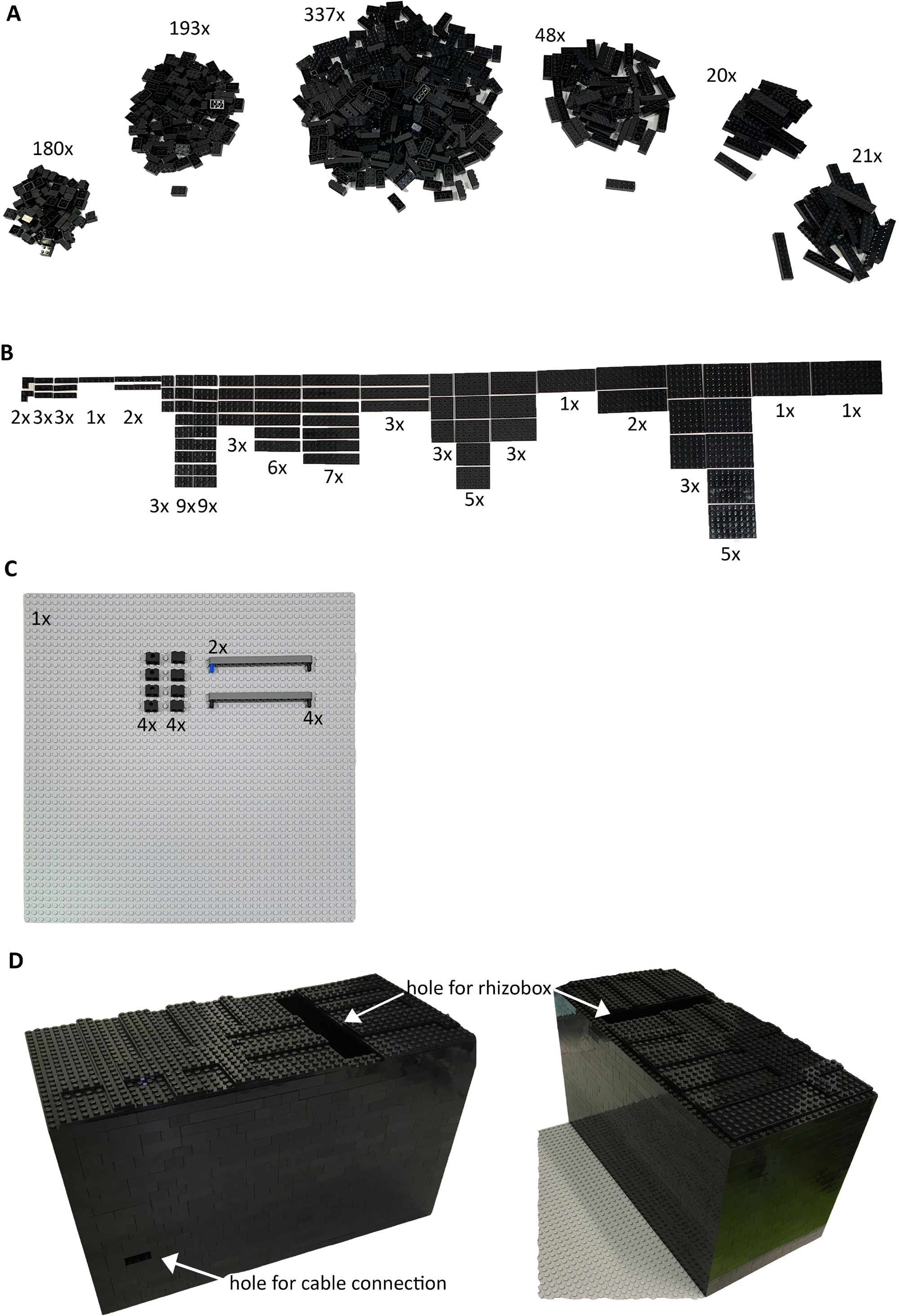
Construction of the BIBLOX. The number of black LEGO® bricks for the construction of the base is indicated (**A**) as well as the number of the plates used as lids (B). One base plate was used and special pieces for the holders (**C**). The finished BIBLOX is in total 22 rows high and includes place for the light source, the camera and the rhizobox (**D**). Note the hole for inserting the rhizobox and one hole for the cables from the power supply and the Raspberry Pi3 system.

The semi-automated root phenotyping method allowed us to analyze the following parameters over time: root area, convex hull area, total root length, and the maximum root system width (**Figures S2A-C**). Additionally, our system allowed us to analyze the angle from the seminal roots (SRGA) described by (Oyanagi et al., 1993; Manschadi et al., 2008) using ImageJ (**Figure S2D**). The comparison of the root morphology of the spring barley trait GP and the facultative trait BCC93 shows a difference in the SRGA at 7 DAS (**Figure 6A**). GP shows a broader angle (mean: 109.3°; n = 7) compared to BCC93 (mean: 62.7°; n = 4). This result underlines the robustness of our system since the SRGA of BCC93 was previously measure with 68.66° (Jia et al., 2019).

**Figure 6.**
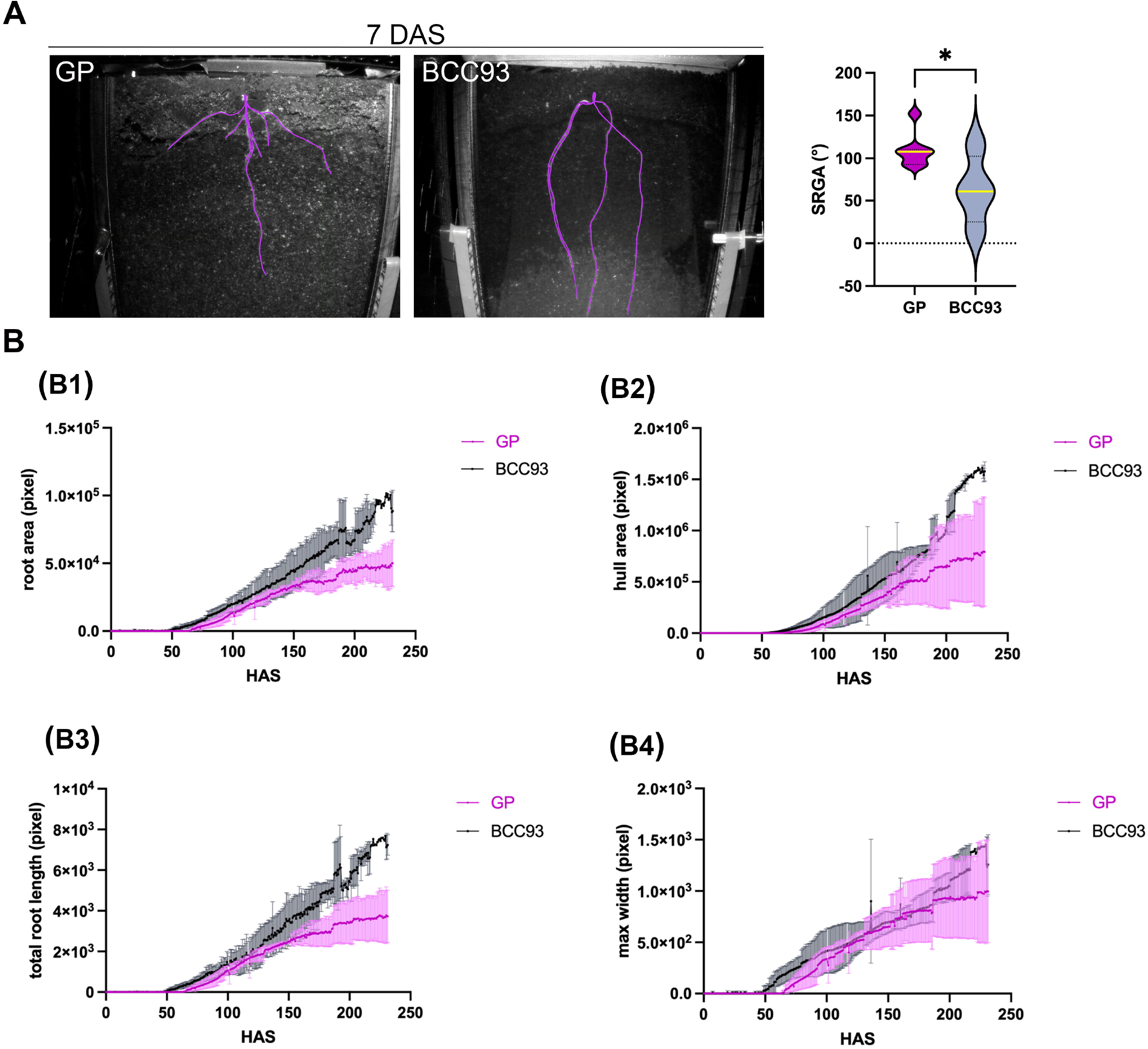
Analysis of root growth and morphology over time. (**A**). Analysis of the seminal root growth angle (SRGA) at 7 DAS. The Violin plot shows a significant difference of the SRGA between GP and BCC93 (n = 4 – 7). * represents ≤ 0.05. (**B**) Root traits measured over time, convex hull area (B1), total root length (B2), maximum root system width (B3), and primary root angle (B4) for three to six biological replicates.

The root tracking shows that the first root could be observed between 64 and 70 hours (3 DAS) and the analyzed parameters resulted in highly reproducible results during early root development until 150 hours (approximately 6 DAS) (**Figure S2B, Figure 6B**). We could observe a difference of the root and hull area, and the total root length of the RSA analysis between GP and BCC93 (**Figure 6B B1, B2, B3**), but no difference in the max root width (**Figure 6B B4**).

## 4. Discussion

Plant performance is strongly affected by environmental conditions. Subsequently, results of controlled conditions of the lab – “Pampered inside, pestered outside” - are often not suitable to translate back to field conditions *(Poorter et al*., *2016)*: *“Besides phenotypically differences between lab- and field-grown plants, the shoot and root environment and the effects of plant density must be considered”*. Thus, the transfer of environmental conditions to controlled lab conditions will obviously improve the knowledge translation gained under lab conditions back to nature.

In nature, roots are growing below the surface in soil. Thus, our first step was the measurement of the environmental parameters in the field to transfer these parameters to controlled lab conditions. Since our research focus is on germination, early root, and grain development of spring barley, we have been measuring soil water content, soil moisture, soil temperature and air temperature in natural fields where spring barley is sown. As we have measurements for three years (Dermendjiev et al., 2021 and 2021, 2022), our data is quite robust and allowed us to set germination and early root development temperature and soil water content at environmental conditions #asnearaspossibletonature.

Roots develop hidden underground in the dark and are only illuminated by the light that penetrates the first 10 millimeters of the soil (Tester and Morris, 1987). Subsequently, experimental conditions in the lab, where roots are often exposure to light, interrupt the root growth development and should be avoided. Within the past decade, RSA traits have been assessed in the lab non-invasively by 2D and 3D imaging techniques (Heeraman et al., 1997; Tracy et al., 2010; Zhu et al., 2011). 3D imaging techniques such as X-ray computed tomography and magnetic resonance imaging have been used to overcome the low spatial resolution often associated with 2D imaging. Whole-plant phenotyping is enabled by phenotyping platforms that allow simultaneous measuring of roots and shoots (Nagel et al., 2012; Jansen et al., 2014). However, high costs of 3D systems (Zhu et al., 2011) and phenotyping platforms still remains.

Rhizoboxes are efficient tools for 2D RSA analysis and enable root development analysis in natural environmental conditions considering parameters like the substrate (e.g. soil), the temperature- and moisture gradient in the soil, the nutrient availability, and the microbiome. Since the first rhizobox, invented in 2008, the construction of rhizoboxes has been optimized and their flexible construction allows the RSA analysis of many different plants, from crops to *Prunus* spp. seedlings (Figueroa-Bustos et al., 2018; Schmidt et al., 2018; Jia et al., 2019; Lesmes-Vesga et al., 2022).

Finally, we emphasized on setting up experiments that include more sustainable research to reduce the anthropogenic climate change. We used secondhand LEGO® bricks and produced 3D-printed rhizoboxes with bio-degradable materials. The usage of local and reusable material enables us to reduce the CO2-footprint in our lab. Of course, this is only the first step, and we have to optimize our experimental setup to further reduce our CO2-footprint to a minimum.

Our focus was to develop a DRD that is especially suitable for the analysis of specific and focused molecular biology-related investigations (e.g. reverse and forward genetic approaches) and for analyses of the RSA in response to distinct environmental factors including different substrate composition. Our set-up allows follow-up molecular, biochemical, -omics and physiological approaches of different crops (**Figure 7**). Since we used LEGO® bricks, our bench-top BIBLOX is flexible regarding its size and is easily relocatable. This flexibility will be extremely helpful for future experiments to investigate and adapt soil temperature and moisture descent-gradient environment for the root (González-García et al., 2022).

**Figure 7.**
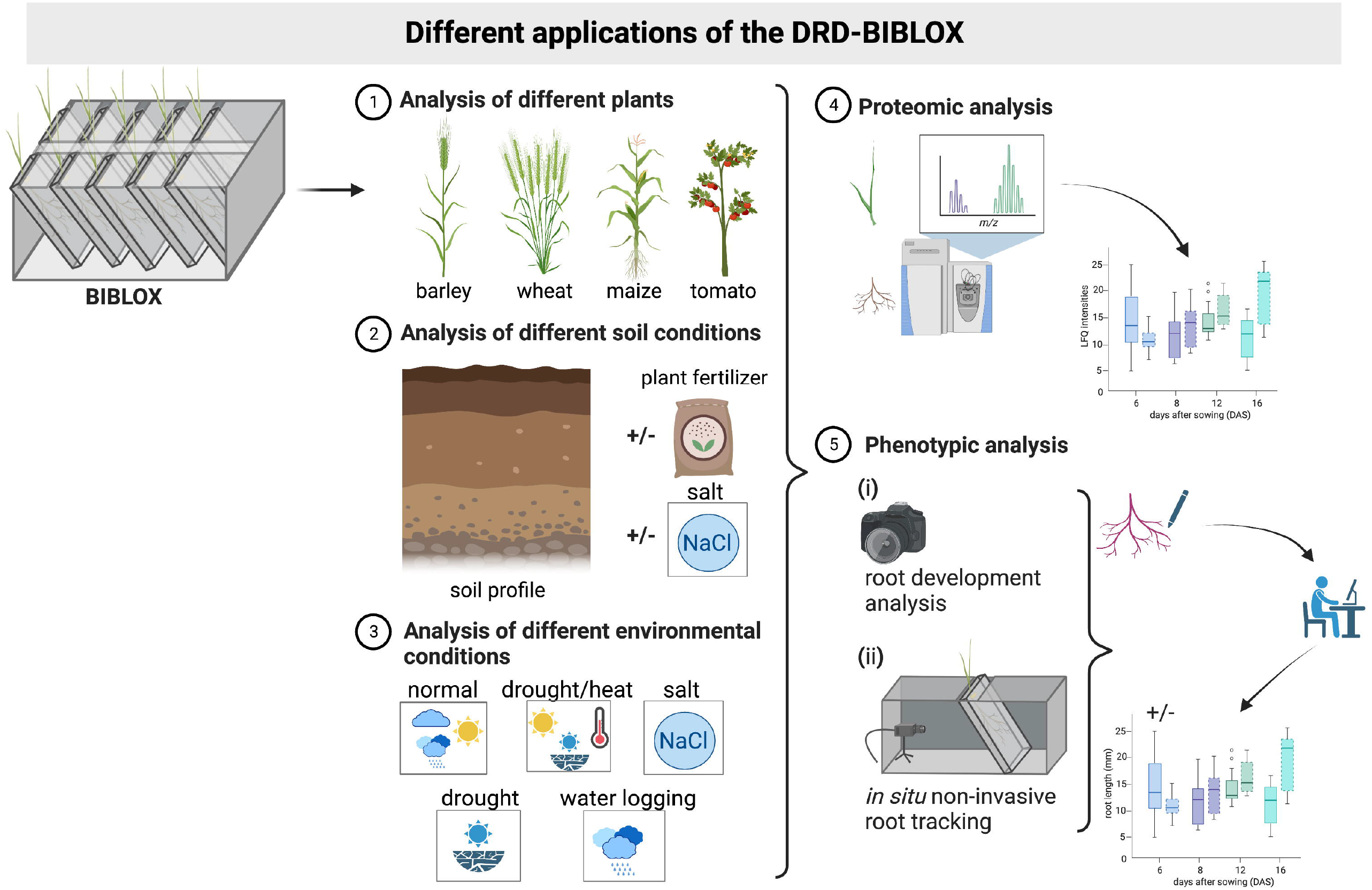
Schematic representation of the different applications of the BIBLOX. Schema was illustrated @Biorender.

## Conclusion

Here, we present our open-hardware tool, the BIBLOX, which is an inexpensive, very flexible, temperature-and humidity resistant DRD, that allows barley germination and root development in soil in the dark with applied environmental parameters mirroring natural environmental conditions (soil temperature, air temperature, soil-moisture). Finally, the BIBLOX provides an imaging application for dark-doot tracking controlled by a Raspberry Pi that enables an easy-to-use, reproducible, inexpensive, and a non-invasive RSA phenotyping approach. Recapitulating, the BIBLOX is a novel system that allows non-invasive *in situ* early root tracking of several crops under controlled environmental conditions #asnearaspossibletonature whilst being #asaccessibleaspossible and #sustainable.

## 6 Data availability statement

The raw data supporting the conclusion of this article will be made available by the authors, without undue reservation.

## 7 Author Contributions

VI, GD, JZ and MS designed the experiment. GD, ES, SP and MS wrote the scripts. ES designed and fabricated the 3D printed rhizobox and performed the image analyses. GD designed the non-invasive root tracking experiment. VI traced the images and performed the SRGA analyses. MS conducted the biological growth analysis in the BIBLOX. TN, GM, and DL performed the LC-MS/MS experiments, MS performed the data analysis. AT harvested, analyzed the bio-organic soil and performed the set-up of the soils #asnearaspossibletonature. SP, MS and VI mounted the data sensors and loggers in the field, SP analyzed the data logger. VI and MS wrote the manuscript. All authors contributed to manuscript preparation and editing.

## 9 Acknowledgments

We thank the master gardeners Thomas Joch and Andreas Schröfl. We would like to thank the head of the Workshop Faculty of Lifescience Heinz Pfeiffer for constructing the PVC rhizoboxes. The authors are thankful to Kerstin Neumann for sharing plant material (barley BCC93). We thank Alois Schweighofer, Lilian Kang and Katarzyna Retzer for critically reading the manuscript. We are thankful to the farmers Leopold Ripfl and Robert Thür for using their fields for data measurement and harvesting field soil. We thank Alexander Seidel for 3D printing.

## 10. Supplemental Material

**Figure S1**. Set up of the field experiment on a barley field measuring environmental parameters. (**A**) A schematic presentation of the field and a picture made by a drone are showing the position of the sensors within the field in lower Austria. (**B**) The measurements of the sensors of the soil temperature and soil moisture in -20 cm depth, the PAR values, and the air temperature. For the soil temperature and moisture only two sensors are presented due to missing measurements of the third sensor, respectively. The black line of the soil moisture graph indicates the mean value.

**Figure S2**. Gallery of images taken between 15 hours and 230 hours of GP DGR in the BIBLOX device at 14°C/12°C day/night cycle in a plant growth chamber. (**A**) Series of images taken at 15 hours intervals (**B**) The same images as in (**A**) traced manually to enhance the contrast between the root and the soil for further semi-automatic analyses. (**C**) Representative output from the image analysis at 76 hours (76 h), 152 hours (152 h) and 230 hours (230 h). The blue line is the maximum root system width and height, the green line is the convex hull and the red shows the analyzed roots. (**D**) Representative image of the SRGA calculation.

**Figure S3**. PCA and loading blot of proteins of 8 DAS. (**A**) PC1 (68.7 %) separates the proteins of shoots and roots, whereas PC2 (17.6 %) separates the proteins depending on the illumination of the root. (**B**) The loading plot indicates the proteins that contributes most to the distribution in PC1 and PC2. **(C)** The contributions plots for PC1 (C1, C2) and PC2 (C3, C4) show how much the total number of proteins (2158) (C1, C3) and the top 30 proteins (C2, C4) contributed to the distribution in PC1 and PC2 respectively. The y axis shows the extent of contribution in %, and the x axis shows the number of the different proteins according to **Supplemental Data 6**.

**Figure S4**. PCA and loading blot of proteins of 16 DAS. (**A**) PC1 (72.5 %) separates the proteins of shoots and roots, whereas PC2 (15.5 %) separates the proteins depending on the illumination of the root. (**B**) The loading plot indicates the proteins that contributes most to the distribution in PC1 and PC2. **(C)** The contributions plots for PC1 (C1, C2) and PC2 (C3, C4) show how much the total number of proteins (2158) (C1, C3) and the top 30 proteins (C2, C4) contributed to the distribution in PC1 and PC2 respectively. The y axis shows the extent of contribution in %, and the x axis shows the number of the different proteins according to **Supplemental Data 6**.

**Figure S5**. PCA and loading blot of proteins of 8 and 16 DAS. (**A**) PC1 (53.5 %) separates the proteins of shoots and roots, whereas PC2 (13.7 %) separates the proteins depending on the illumination of the root. (**B**) The loading plot indicates the proteins that contributes most to the distribution in PC1 and PC2. **(C)** The contributions plots for PC1 (C1, C2) and PC2 (C3, C4) show how much the total number of proteins (2158) (C1, C3) and the top 30 proteins (C2, C4) contributed to the distribution in PC1 and PC2 respectively. The y axis shows the extent of contribution in %, and the x axis shows the number of the different proteins according to **Supplemental Data 6**.

**Figure S6. Test arrangement of several BIBLOXes in one growth chamber**. Note the experiment in the middle where the roots are not protected from light.

**Supplemental Data 1**. R-script for data logger analysis.

**Supplemental Data 2**. 3D-CAD software Fusion 360 providing data for rhizobox.

**Supplemental Data 3**. Official report of the tested bio-organic field soil. **Supplemental Data 4**. Official report of the tested conventional field soil. **Supplemental Data 5**. Excel sheet of proteomic analysis results.

**Supplemental Data 6**. Excel sheet of proteomic analysis with data used for analysis in R (PCA).

**Supplemental Data 7**. R code used for analysis of proteomic analysis results

**Supplemental Video 1**. Drone video of the barley field with installed loggers and sensors.

**Supplemental Video 2**. Construction of the BIBLOX for root tracking.

**Supplemental Video 3**. Root tracking of GP. Pictures were taken all 4 minutes in the period of 230 hours.

**Supplemental Video 4**. Root tracking of BCC93 over 192 hours. Pictures were taken all 4 minutes in the period of 192 hours.

