## Supplementary figures and images for "A bench-top dark-root device built with LEGO® bricks enables a non-invasive plant root development analysis in soil conditions mirroring nature"

### Figure S1

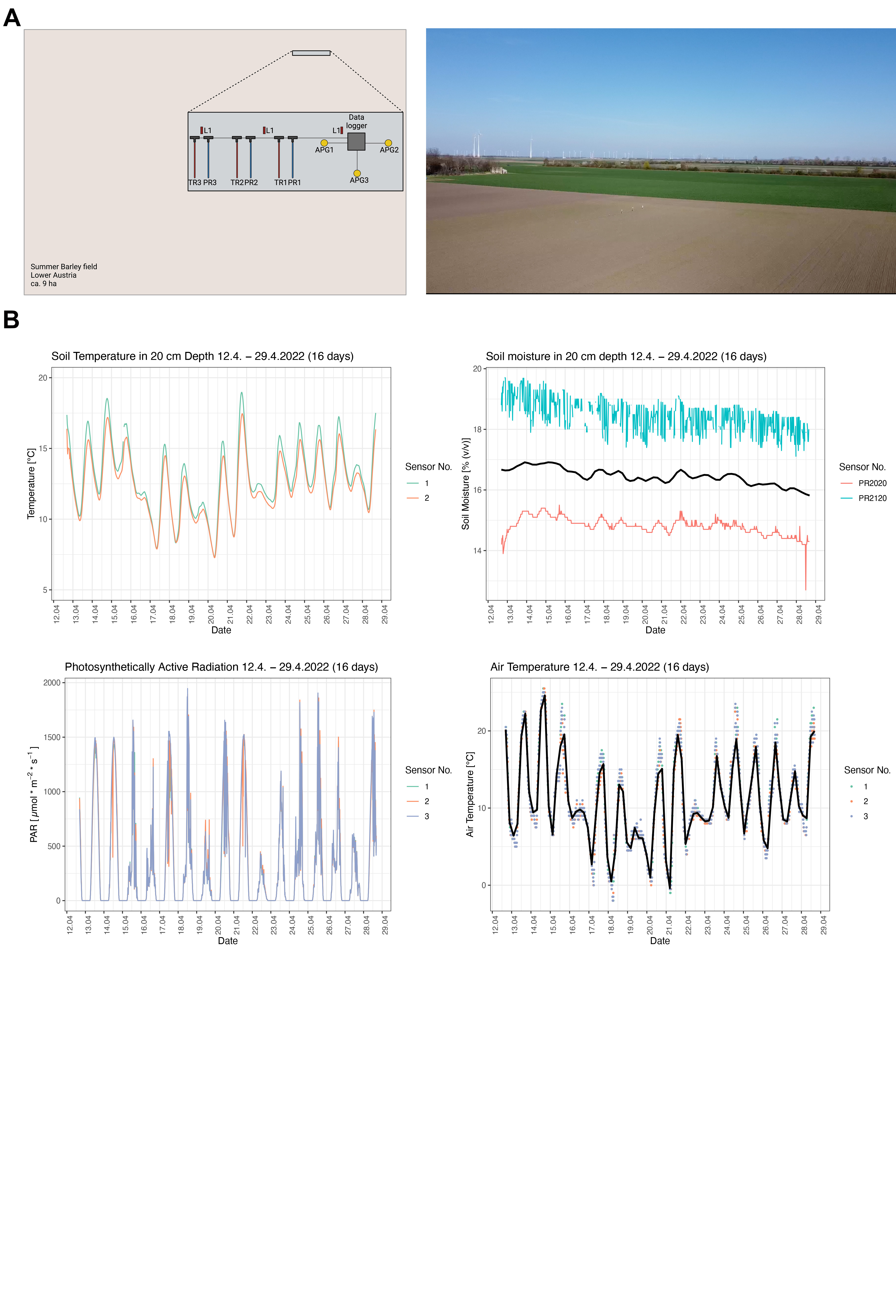

### Figure S2

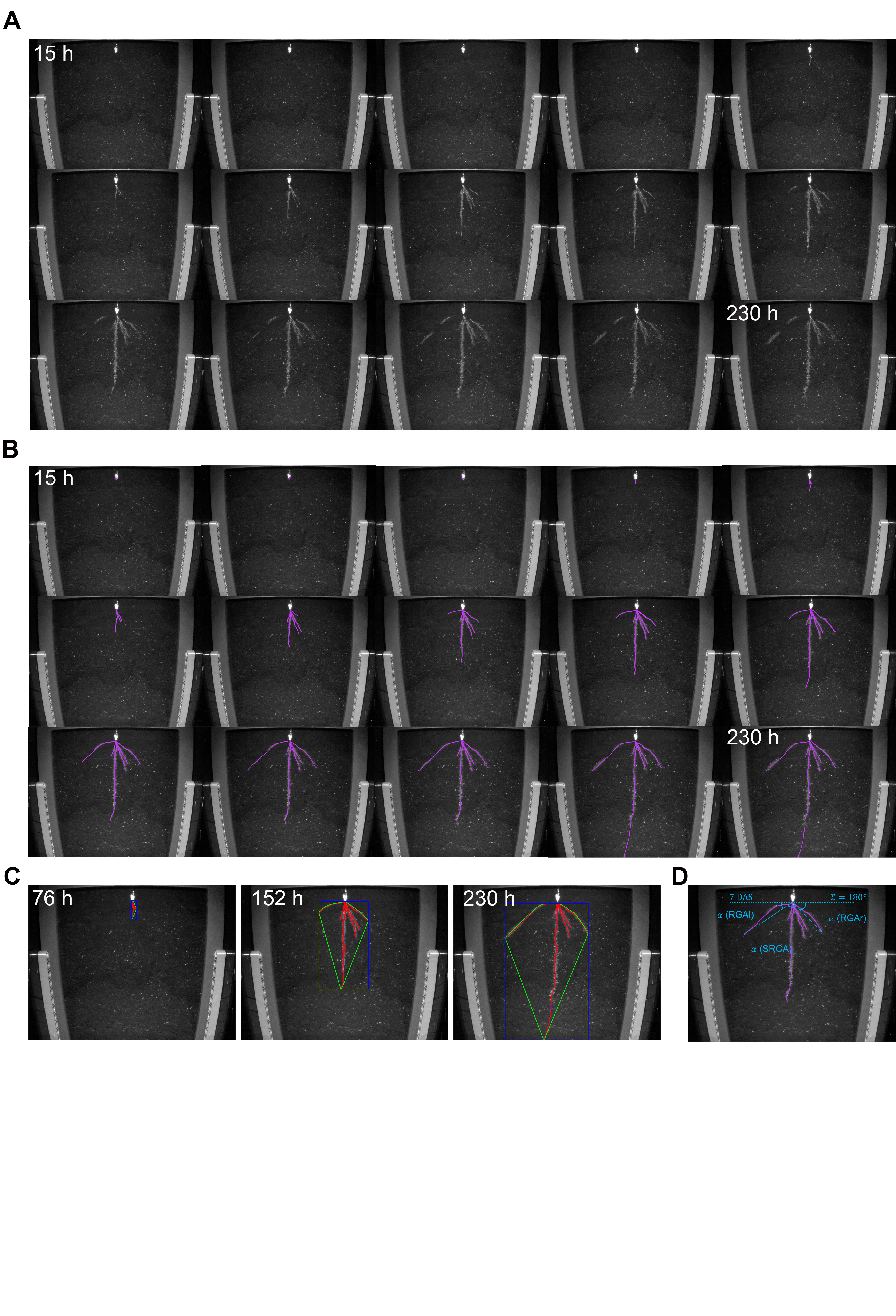

### Figure S3

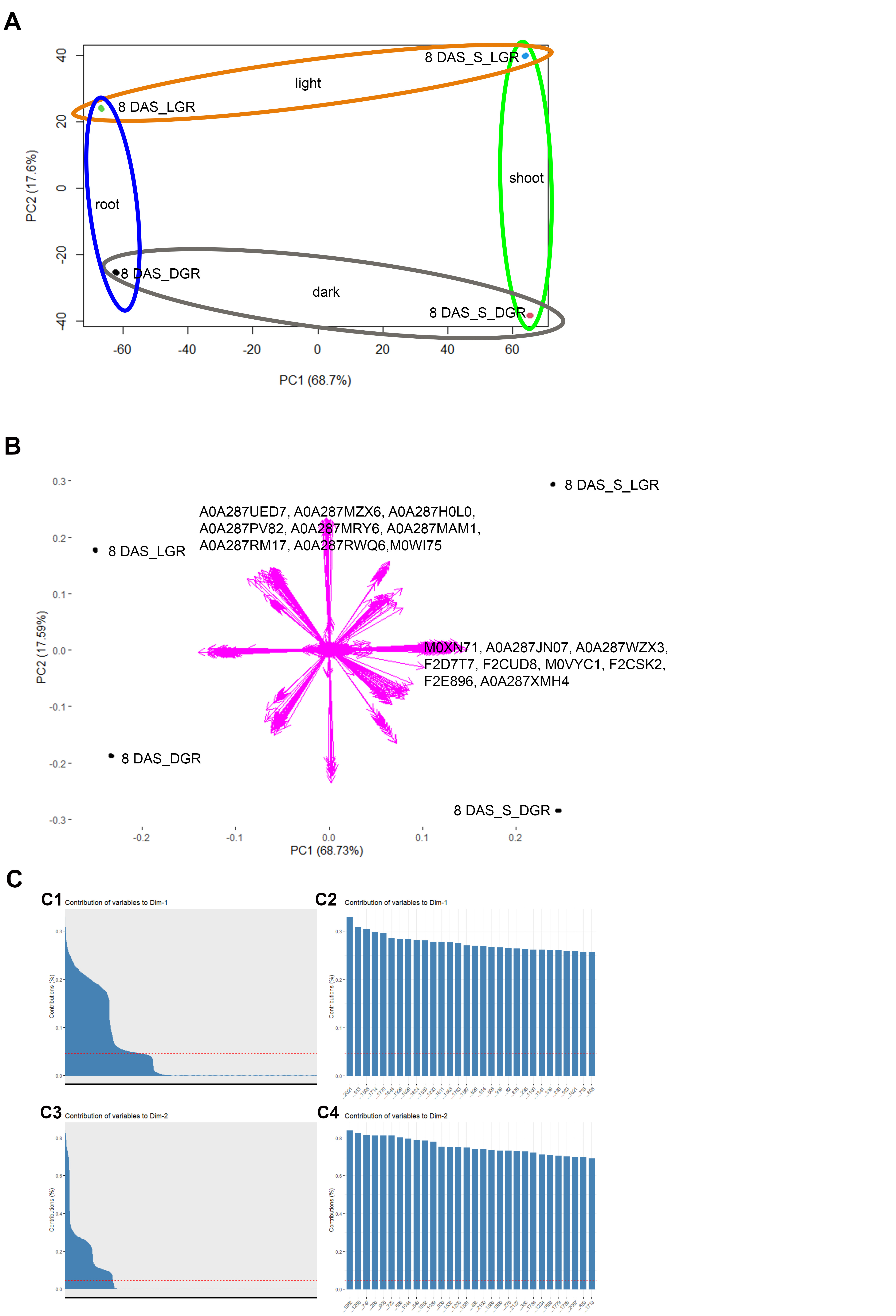

### Figure S4

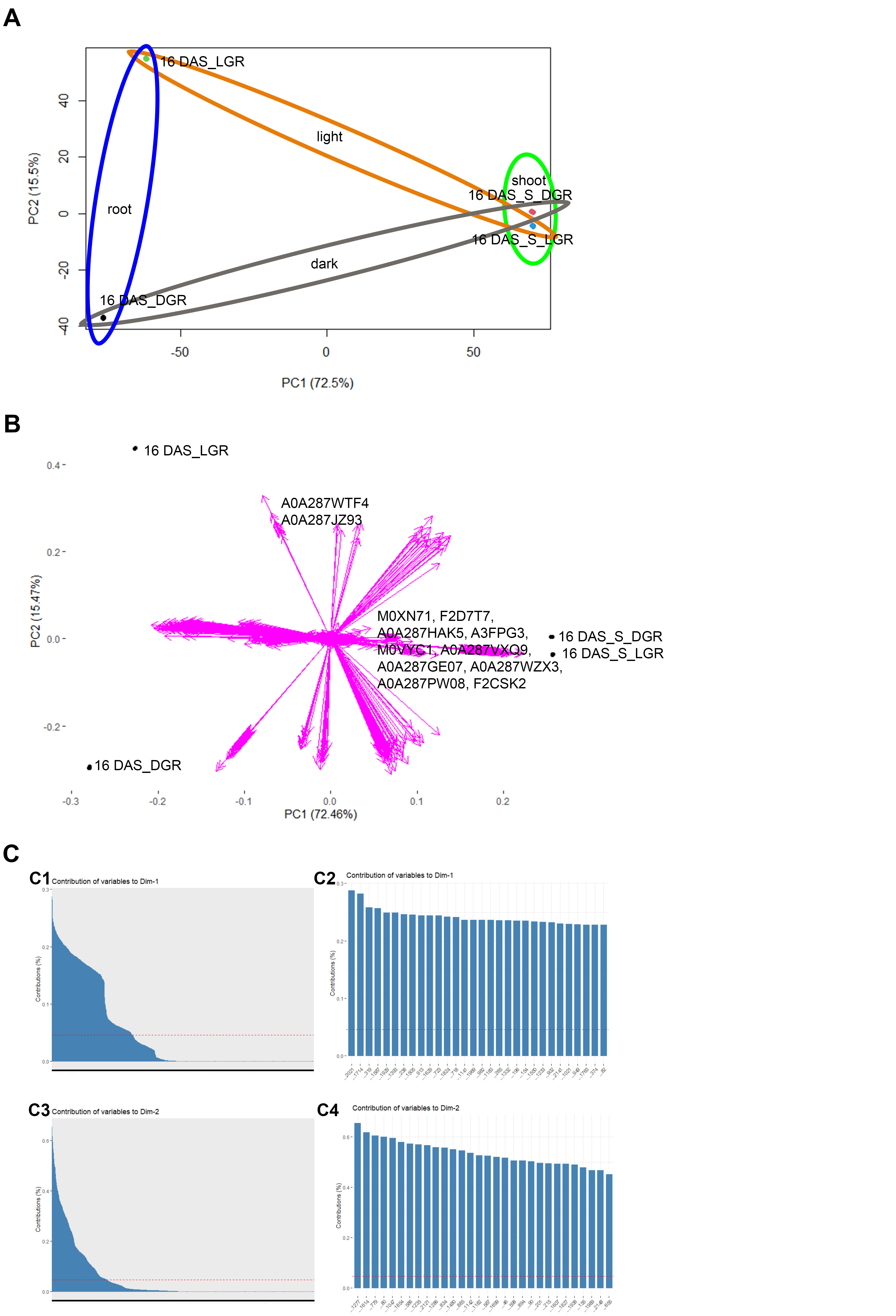

### Figure S5

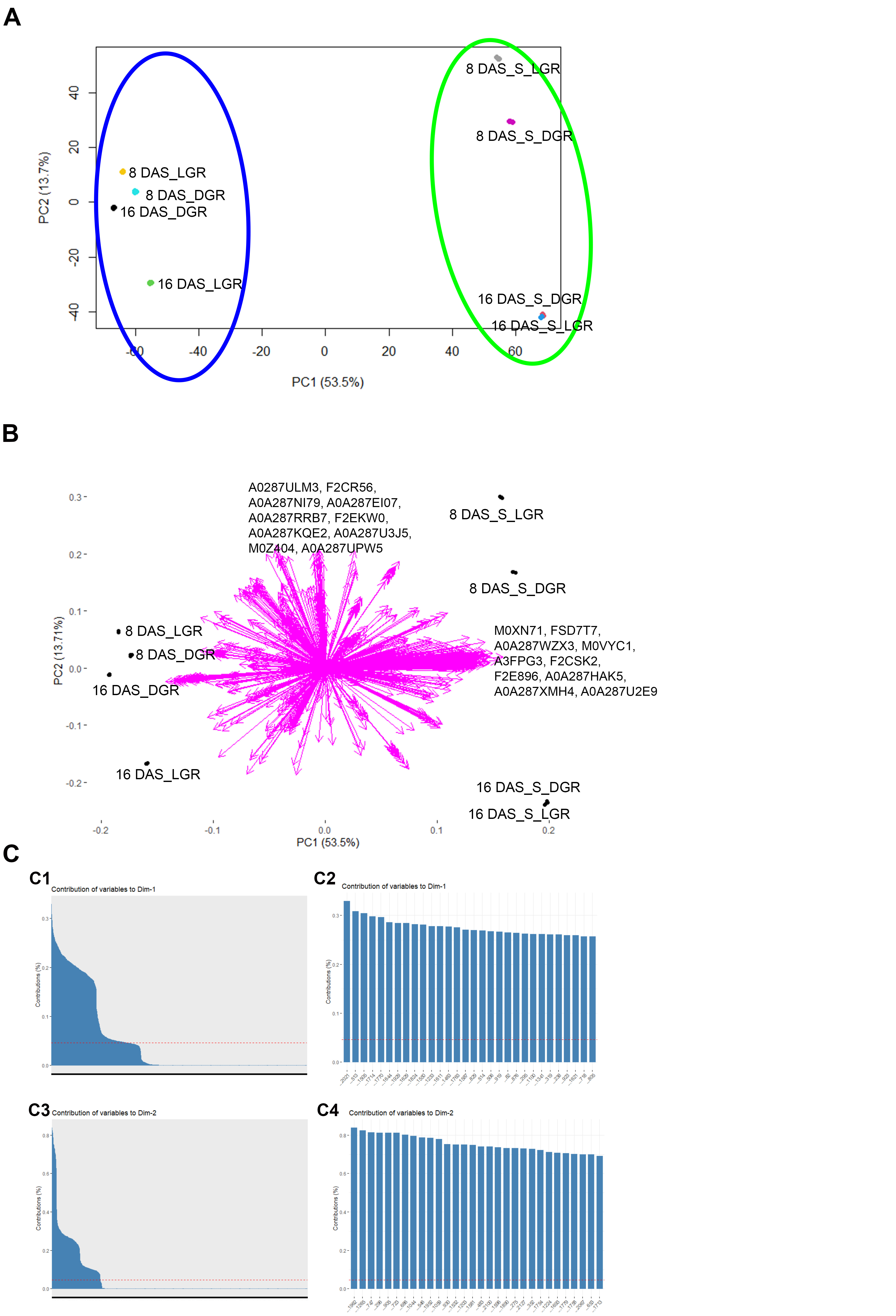

### Figure S6

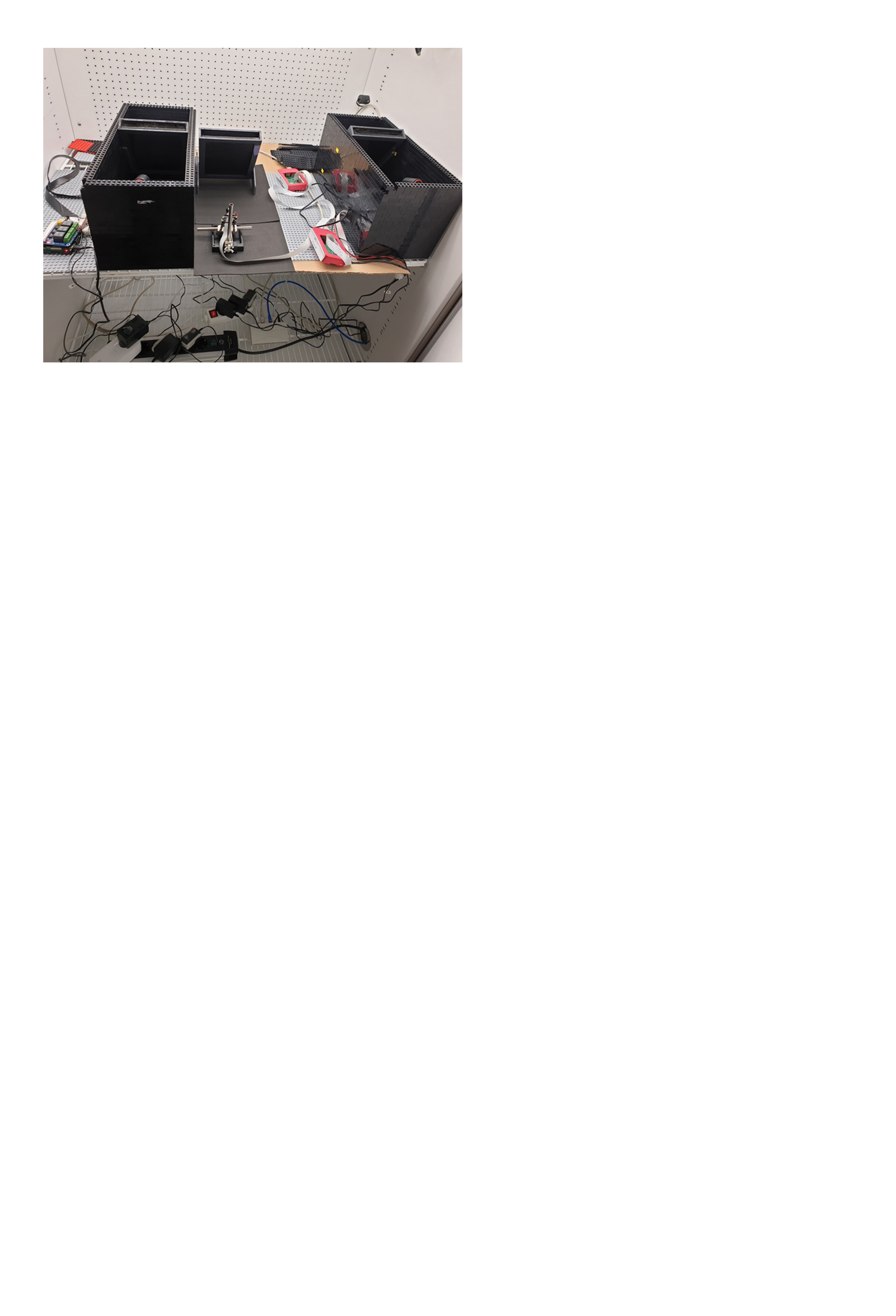
